# A Microfluidic Device to Simulate the Impact of Gut Microbiome in Cancer

**DOI:** 10.1101/2022.11.13.516284

**Authors:** Ekansh Mittal, Youngbok (Abraham) Kang

## Abstract

The gut microbiome has a role in the growth of many diseases such as cancer due to increased inflammation. There is an unmet need to identify novel strategies to investigate the effect of inflammation mediated by gut microbiome on cancer cells. However, there are limited biomimetic co-culture systems that allow to test causal relationship of microbiome on cancer cells. Here we developed a microfluidic chip that can simulate the interaction of the gut microbiome and cancer cells to test the effects of bacteria and inflammatory stress on cancer cells in vitro. To quantify the effect of bacteria on the growth of colorectal cancer cells, we cultured colorectal cancer cell line with Bacillus or lipopolysaccharide (LPS), which is a purified bacterial membrane and induce major inflammatory response, in the PDMS microfluidic device. We found that both LPS and Bacillus significantly accelerate the growth of colorectal cancer cells. These results show that the increased presence of certain bacteria can promote cancer cell growth and that these microfluidic chips can be used to test the specific correlation between bacteria and cancer cell growth. These microfluidic devices can have future implications for various cancer types and to identify treatment strategies.

## 1. Introduction

Emerging studies suggest that the gut microbiome may have a role in cancer initiation and progression [1- 3]. The microbiome consists of trillions of good and bad microorganisms that colonize in our body, and about 70% of those reside in our gut. These bacteria play a vital role in maintaining homeostasis in the body. An imbalance in bacterial composition may cause diseases including cancer [4, 5] that might be due to increase in inflammation [6]. Most of these studies have shown the association of gut microbiome with increased cancer progression; however, limited studies have established a causal relationship [2, 7].

Therefore, it is important to develop a novel strategy to investigate the relationship between the gut microbiome and cancer cells. In our study, we tested the effect of gut microbiome on the cancer cell growth in the microfluidic device.

## 2. Materials and Methods

### 2.1. Fabrication of the microfluidic device

The device consists of one middle chamber (width (0.8 cm) x length (2 cm) x height (250 µm)) for cancer cell culture and two side channels (width (0.1 cm) x length (1.5 cm) x height (250 µm)) to introduce media and bacteria. The template of the microfluidic device was made of SU-8 photoresist of 250 μm thickness on a silicon wafer using a photolithography technology. The polydimethylsiloxane (PDMS) devices were molded from the SU-8 template according to soft-lithography process. [8] The PDMS microfluidic devices were bonded to a glass slide after air plasma treatment. The volume of the cell culture chamber in a microfluidic chip was 160 µL (250 µm height of micro channel). Later, to improve efficiency of the device operation and simplify a fabrication of a device, we 3D-printed the template of the device.

### 2.2. Co-culture of cancer cells and bacterial cells

A human colorectal carcinoma cell line (red fluorescent protein (RFP)+ HCT116) was seeded into the cell culture chamber of the device and cultured in McCoy’s 5a medium containing 10% FBS at 37^0^C and 5% CO2. Media was fed at 5 µL/min through side channels by a syringe pump (World precision instruments). Once cells make a confluent monolayer, bacterial cells were introduced through the side channel, diffused to a cell culture chamber, and cultured at the same condition with cancer cell culture.

### 2.3. Quantification

Zeiss Fluorescent microscope was used to record all the images at 20X resolution. Image J was used to quantify the individual cells from 4 areas for each chip in duplicate. The data is represented as an average with standard error. Paired Student t-Test was used to calculate the significance.

## 3. Results

To test the causal effect of bacteria on cancer progression, we developed a simple microfluidic device for simulating the conditions of the gut. A PDMS microfluidic device has three inlets for injecting cells and media and outlets for waste collection. We first coated the cell culture chamber of a device with collagen (Corning # 354249) to provide an extracellular matrix. Then we seeded 40,000 colorectal cancer cells (HCT-116) in the middle cell culture chamber. The media from the side chambers then flows into the middle chamber to make a confluent cell monolayer. To replicate tumor hypoxic microenvironment, we used the media supplemented with concentration of 1% sulfite and 100 µM cobalt for removal of oxygen and acceleration of oxygen depletion. According to our previous data, hypoxic environment (less than 1% oxygen) is generated in the cell culture chamber using this method [9].

When cells are grown in the hypoxic environment, about 1,000 Bacillus bacteria cells were loaded into the side chambers. Bacillus was used due to BSL-1 level. The chip also allowed to maintain liquid flow during co-culturing of bacterial and cancer cells using syringe pumps. 10 ng/ml lipopolysaccharide(LPS) was also used as a positive control. LPS is a purified bacterial membrane and known to induce inflammatory response [10]. We quantified cell morphology and cell growth over time through a fluorescent microscope.

We found that both LPS and Bacillus accelerated the growth of cancer cells compared to vehicle treated cancer cells over time. After 4 days of culturing, LPS promoted the growth of cancer cells 2.02 fold (p value = 0.012) while stimulation of bacterial cells increased the growth1.58 fold (p value = 0.011) as compared to vehicle treated cells (Figure 1B-C). These results show that increased presence of certain bacteria or microbial inflammatory stress can promote colorectal cancer cell growth. Further, the microfluidic chip can be used to test the causal relation between bacteria and cancer cell growth.

**Figure 1.**
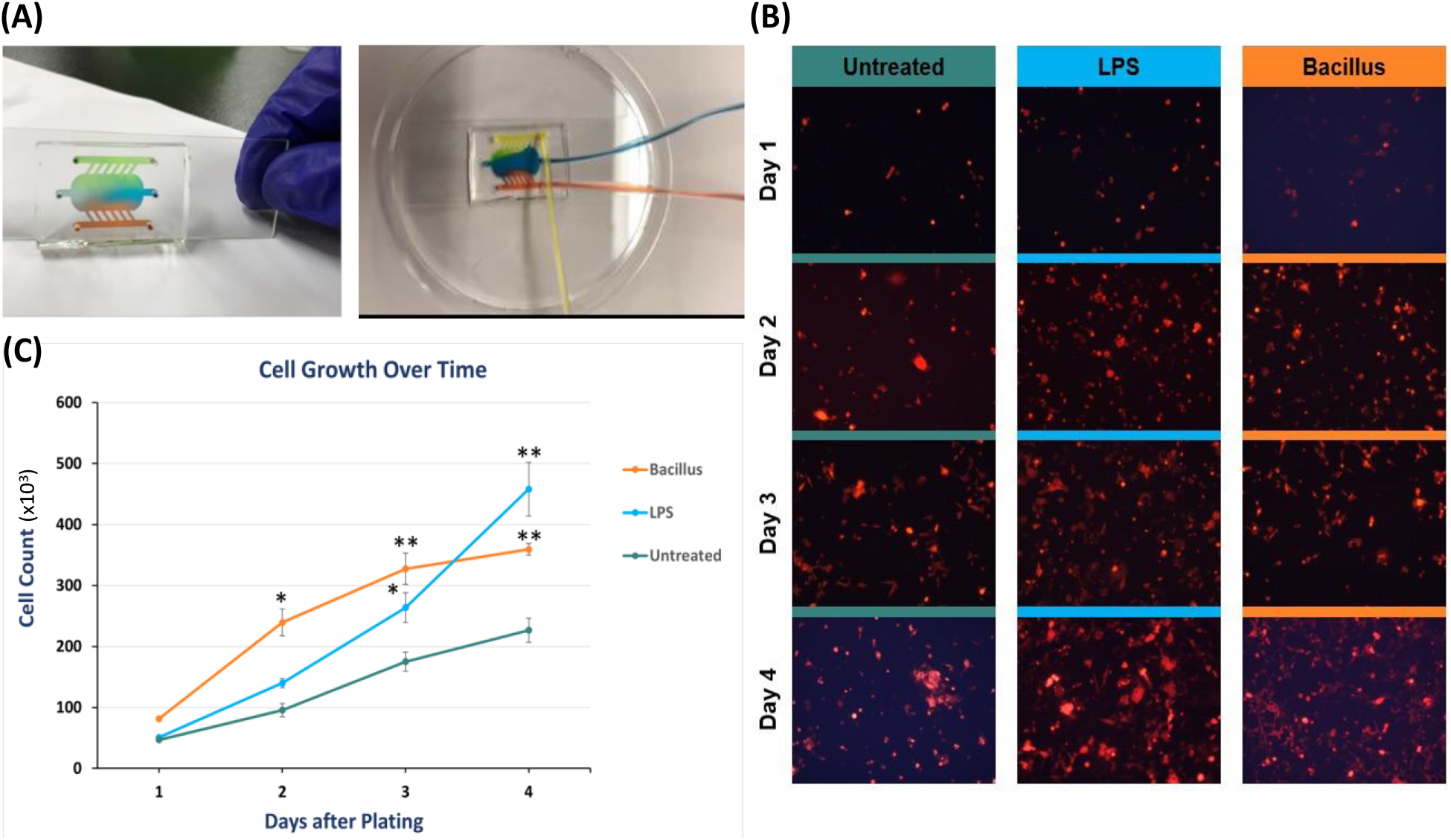
The abundance of gut bacteria promotes the growth of colorectal cancer cells over time. (A) The microfluidics design that allow bacteria and cancer cells to co-culture. In outer inlets total 1000 bacterial cells were loaded. Middle inlet was used to inject 40,000 RFP labeled HCT 116 cells. Cells were cultured for 24 hrs and then microchannels were washed thoroughly to washout non-adherent HCT 116 cells and extra bacterial cells (B, C). Then microfluidics was attached to syringe pump that injected media at 5 ul/min flow rate. The effect of Bacillus bacteria or lipopolysaccharide (LPS; 10ng/ml) on the growth of red fluorescent labelled colorectal cancer cells (HCT 116) over time that is quantified daily by imaging 4 independent areas. The images were counted using Image J analysis. Student t-test was used to calculate p Value. * < 0.05 and ** <0.01.

## 4. Discussion

Using a microfluidic device, we demonstrated that the presence or abundance of bacteria can significantly promote the growth of cancer cells. There are several possible mechanisms by which imbalance in bacteria community can impact cancer cell growth. Almost all bacteria secrete substances such as lipopolysaccharide (LPS) and peptidoglycan, which are found in the bacterial cell membrane [10]. When these substances enter the gut and bind to the receptors on the myeloid cells found in the gut. These interactions cause epigenetic reprogramming leading to change in gene expression profile and lead to increased pro-inflammatory cytokine secretion, reduce the clearance of pathogens and cancer cells, reduce cancer cell killing, and reduce intestinal barrier fortification, allowing more pathogens to enter [2].

LPS and peptidoglycan also increase the secretion of pro-inflammatory cytokines, increasing cancer progression. When these compounds bind the toll-like receptor (TLR2, TLR4) on the cell membrane, the receptor signals to the IRAK kinase, which signals to the NF-kB1 transcription factor [10] with increased secretion of the proinflammatory cytokines such as IL6 [6]. This pathway can be targeted by drugs such as IRAK-1 kinase and NF-kB1 inhibitors or IL-6 blocking antibody to reduce cancer progression [6]. This microfluidic device has broader implications including simulating other cancer types. This device can test both the effect of bacteria and new treatment on clinical samples for the identification of personalized therapy, thus reducing the need for mouse model for preclinical testing, which is a lengthy and expensive process. Overall, we established a new co-culture system for bacteria and cancer cells and identified that gut microbiome can promote can cell growth. Thus, can serve as a new therapeutic entity.

